# METTL4 catalyzes m6Am methylation in *U2 snRNA* to regulate pre-mRNA splicing

**DOI:** 10.1101/2020.01.24.917575

**Authors:** Yeek Teck Goh, Casslynn W. Q. Koh, Donald Yuhui Sim, Xavier Roca, W. S. Sho Goh

## Abstract

*N*^6^-methylation of 2’-O-methyladenosine (Am) in RNA occurs in eukaryotic cells to generate *N*^6^,2’-O-dimethyladenosine (m6Am). Identification of the methyltransferase responsible for m6Am catalysis has accelerated studies on the function of m6Am in RNA processing. While m6Am is generally found in the first transcribed nucleotide of mRNAs, the modification is also found internally within *U2 snRNA*. However, the writer required for catalyzing internal m6Am formation had remained elusive. By sequencing transcriptome-wide RNA methylation at single-base-resolution, we identified human METTL4 as the writer that directly methylates Am at *U2 snRNA* position 30 into m6Am. We found that METTL4 localizes to the nucleus and its conserved methyltransferase catalytic site is required for *U2 snRNA* methylation. By sequencing human cells with overexpressed *Mettl4*, we determined METTL4’s *in vivo* target RNA motif specificity. In the absence of *Mettl4* in human cells, *U2 snRNA* lacks m6Am thereby affecting a subset of splicing events that exhibit specific features such as overall 3’ splice-site weakness with certain motif positions more affected than others. This study establishes that METTL4 methylation of *U2 snRNA* regulates splicing of specific pre-mRNA transcripts.

## Introduction

*N*^6^-methylation of deoxyadenosine affects the biophysical properties of the modified DNA and also regulates genomic processes (1, 2). Likewise, *N*^6^-methylation of adenosine generates RNA modifications that regulate the expression and metabolism of the host RNA without changing the underlying sequence (3). One resultant RNA modification is *N*^6^-methyladenosine (m6A) that is found internally within various RNA species including mRNAs and rRNAs (4-7). Each of the several identified m6A methyltransferases (writers) that catalyze *N*^6^-methylation of adenosine have different target sequence specificities. For example, METTL3 catalyzes *N*^6^-methylation of DRACH to give DRm6ACH (D=A/G/U, R=A/G, H=A/C/U) (4, 5, 8, 9). METTL3 also functions in a complex with co-factors including METTL14, WTAP, KIAA1429 and RBM15/RBM15B (10-14). This complex mostly targets mRNAs in the region proximal to the stop codon. Another m6A writer is METTL16 that methylates ‘UACAGAGAA’ to give ‘UACm6AGAGAA’. Besides U6 snRNA, METTL16 also targets the *Mat2a* 3’UTR (15). Finally, ZCCHC4 and METTL5 have been identified as the writers responsible for m6A catalysis in 18S rRNA and 28S rRNA respectively (16-18).

Adenosine *N*^6^-methylation of 2’-O-methyladenosine (Am) results in the RNA modification *N*^6^,2’-O-dimethyladenosine (m6Am) that is found in the first transcribed nucleotide adjacent to the RNA methylguanosine cap (Figure S1A) (19, 20). Given its predicted *N*^6^-methyladenine methyltransferase domain and its interaction with the phosphorylated C-terminal tail of RNA polymerase II during RNA transcription, PCIF1 was a strong candidate as a *N*^6^-methyladenine writer (21, 22). Subsequently, PCIF1 was established as the writer responsible for the *N*^6^-methylation of Am at the transcriptional start site (TSS) to generate m6Am (23-27). This facilitated the characterization of TSS-associated m6Am as a regulator of mRNA resistance to DCP2-mediated decapping, and potentially mRNA translation and cell growth (19, 23, 25, 26).

Besides TSS-associated m6Am, m6Am sites have also been mapped to internal RNA sites, specifically at *U2 snRNA* position 30 (20, 28). The functional importance of m6Am in *U2 snRNA* was not well-characterized, likely because the writer responsible for methylating *U2 snRNA* Am30 was previously unknown. In our work here, we utilize a primarily sequencing approach to establish methyltransferase-like 4 (*Mettl4*) as the *U2 snRNA* m6Am writer. By overexpressing *Mettl4*, we uncovered both the *in vivo* target sequence preference of METTL4 and its ability to catalyze m6Am formation internally within mRNAs. Finally, by knocking out *Mettl4* from cells, we demonstrate that the resulting loss of *U2 snRNA* m6Am perturbs splicing of target mRNAs with distinctive features such as weak 3’ splice sites.

## Results

### *U2 snRNA* m6Am is absent in *Mettl4*-KO cells

*Mettl4* is one of a diverse family of proteins that are homologous to the MT-A70 subunit of the m6A writer *Mettl3* (29). In order to assess *Mettl4* as a potential writer, we performed m6A-Crosslinking-Exonuclease-sequencing (m6ACE-seq), which is capable of quantitatively mapping precise *N*^6^-methyladenine methylomes in the form of m6A or m6Am (Figure S1A) (27). Comparison of the RNA methylomes between wildtype (WT) and *Mettl4*-KO HEK293T cells revealed a single site that exhibited almost 70 fold reduced relative methylation level (RML) in the absence of METTL4 (Figure 1A,1B). The single-base-resolution of m6ACE-seq allowed us to map this site to position 30 of *U2 snRNA* (Figure 1C). To verify that this METTL4-dependent adenosine methylation is located within *U2 snRNA*, we isolated various snRNAs for anti-m6A dot blotting (Figure S1B). As expected, we detected an RNA methylation signal specific only to WT *U2 snRNA* but not in *Mettl4*-KO *U2 snRNA* or any *U1* snRNA (Figure 1D).

**Figure 1.**
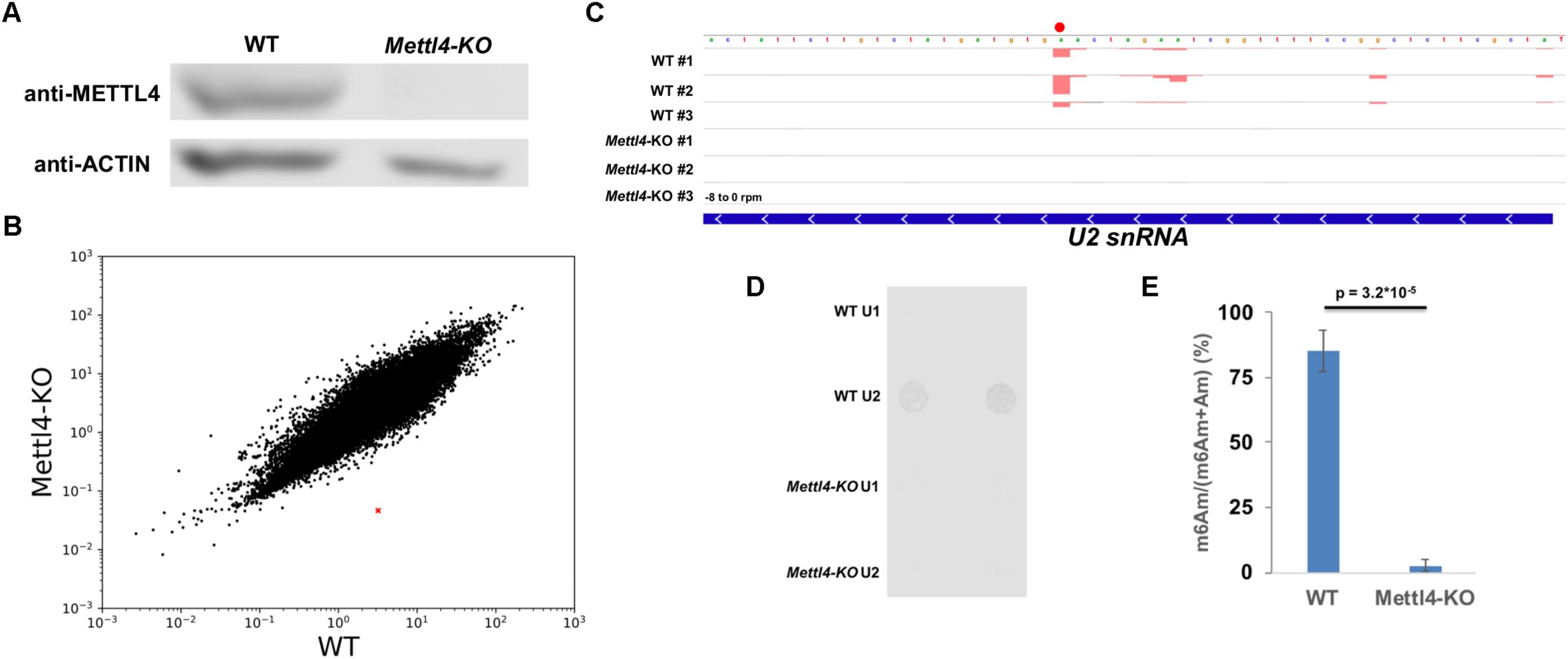
METTL4 mediates m6Am modification in *U2 snRNA*. (A) Western blotting of human METTL4 in WT versus *Mettl4*-KO lysate with ACTIN as a loading control. (B) Scatterplot of average RML of WT versus *Mettl4*-KO cells. *U2 snRNA* is denoted with a red ‘X’. (C) m6ACE (red) and Input (black) read-start counts (in reads per million mapped or RPM) mapped to *U2 snRNA. U2 snRNA* m6Am position identified in (B) is denoted by a red dot. Sequence corresponds to the same strand as the m6Am site. Blue horizontal bar represents transcript. (D) Anti-m6A dot blotting of various snRNAs purified from WT versus *Mettl4*-KO RNA. Duplicate dots are shown. (E) Nucleoside HPLC-MS/MS of m6Am as a percentage of total m6Am and Am in *U2 snRNA* purified from WT versus *Mettl4*-KO RNA. Displayed are average and standard deviation error of biological triplicates. 1-tailed Student’s T-test p-value is shown.

Since the anti-m6A antibody is not able to distinguish between the structurally similar m6A and m6Am modifications (Figure S1A), isolated *U2 snRNA* was digested and subjected to nucleoside high performance liquid chromatography coupled with tandem mass spectrometry (HPLC-MS/MS) to verify the identity of the RNA modification within *U2 snRNA*. While there were no appreciable m6A levels in *U2 snRNA*, we instead found that the majority of Am are in the *N*^6-^methylated m6Am form (Figure 1E, S1C, S1D). More importantly, m6Am was not detected in *Mettl4*-KO *U2 snRNA* (Figure 1E, S1C). We note that the HPLC-MS/MS m6Am signal was derived internally from *U2 snRNA* as we did not use any decapping enzyme in digesting *U2 snRNA*, which would prevent the quantification of any cap-adjacent RNA nucleotide. Furthermore, previous and current RNA methylomes have demonstrated that *U2 snRNA* lacks a TSS-associated adenosine *N*^6^-methylation (Figure 1C) (27). Altogether, this demonstrates that METTL4 is necessary for m6Am formation within *U2 snRNA*.

### *N*^6^-methylation of *U2 snRNA*-internal Am requires the METTL4 DPPW catalytic motif

METTL4 can either directly catalyze the *N*^6^-methylation of Am or act as an indirect co-factor to the actual catalytic writer. To investigate METTL4’s role in RNA methylation, we expressed and purified recombinant C-terminal-3x-FLAG-tagged human METTL4 (METTL4^WT^) (Figure 2A, S2A). Incubation of METTL4^WT^ with a RNA oligonucleotide matching the 35nt 5’ end of *U2 snRNA* (*U2*-Am30, Figure 2B, Table 1) generated a *N*^6^-methyladenosine signal detectable via anti-m6A dot blotting (Figure 2C). Using HPLC-MS/MS, we verified this methylated signal as m6Am and not m6A (Figure 2D). Additionally, METTL4 contains a ‘DPPW’ motif, which are catalytic residues required for methyltransferase activity (29). Therefore, we mutated the METTL4 ‘DPPW’ to ‘APPA’ to generate a catalytically-dead METTL4 (METTL4^CD^, Figure 2A, S2A). Using the same *in vitro* methylation assay, we found that loss of the ‘DPPW’ catalytic site in METTL4^CD^ abrogates its ability to methylate RNA Am into m6Am (Figure 2C, 2D). This demonstrates that human METTL4 directly catalyzes the *N*^6^-methylation of Am to m6Am *in vitro*.

**Figure 2.**
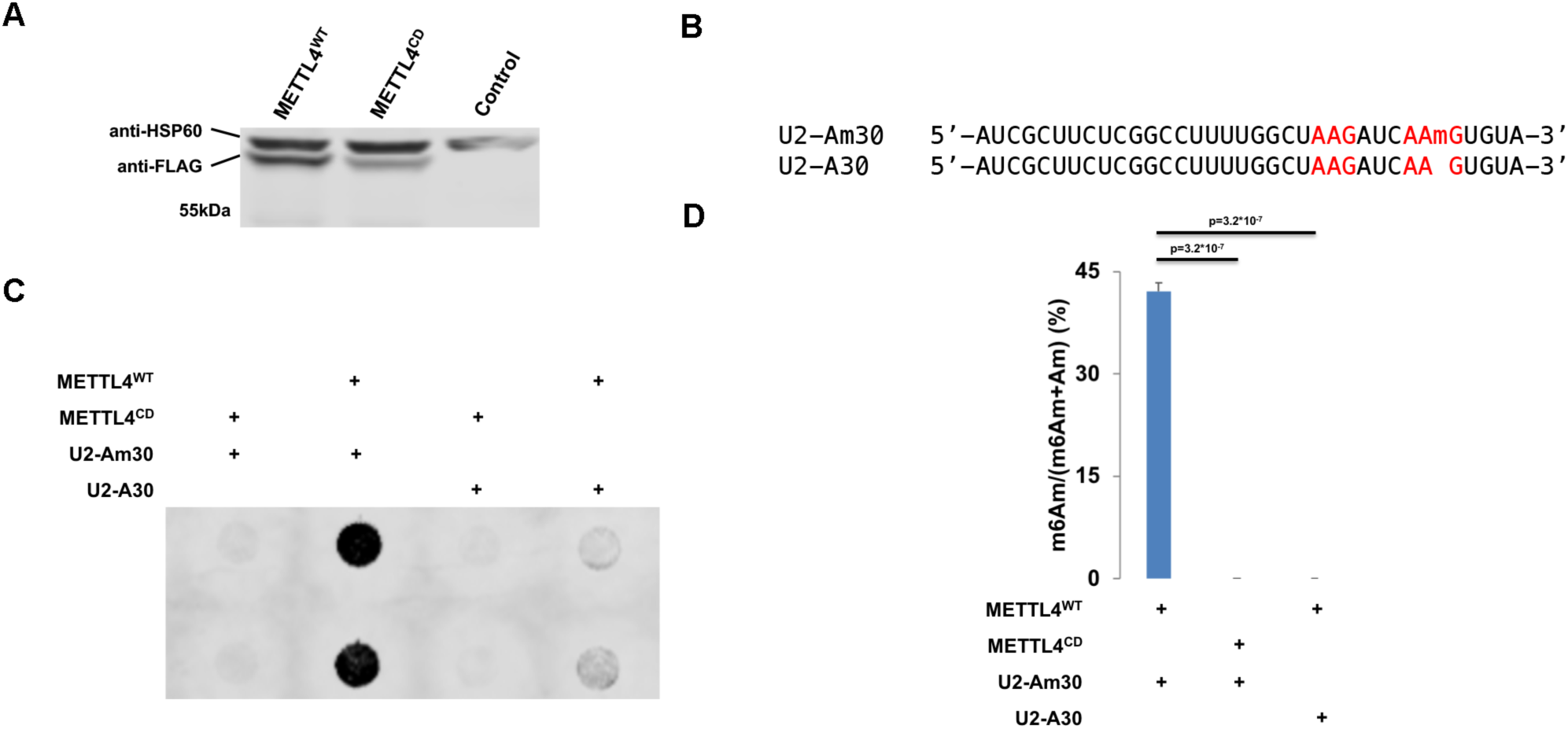
METTL4 directly catalyzes m6Am formation *in vitro*. (A) Western blotting of cell lysate from *Mettl4*-KO HEK293T transfected with METTL4^WT^ or METTL4^CD^ or untransfected (control). Expected size of purified protein is 57kDa. HSP60 (∼61kDa) is displayed as a comparison. (B) Sequence of *U2*-Am30 and *U2*-A30 RNA substrates used for *in vitro* methylation. ‘AAG’ sites are highlighted in red. (C) Anti-m6A dot blotting of RNA substrates *in vitro* methylated by METTL4^WT^ or METTL4^CD^. Duplicate dots are shown. (D) Nucleoside HPLC-MS/MS of m6Am as a percentage of total m6Am and Am in RNA substrates *in vitro* methylated by METTL4^WT^ or METTL4^CD^. Displayed are average and standard deviation error of triplicates. 1-tailed Student’s T-test p-value is shown.

Given the structural similarity between Am and A, we next tested if METTL4 is also able to methylate A into m6A (Figure S1A). We subjected a similar *U2 snRNA*-based RNA substrate with the single Am replaced with A (*U2*-A30, Table 1) to *in vitro* methylation by METTL4^WT^ (Figure 2B). This led to quantifiable m6A signals, albeit at a lower level than that of METTL4^WT^ *in vitro* methylation of *U2*-Am30 (Figure S2B). Expectedly, this methylation activity was absent for METTL4^CD^. Both *U2*-A30 and *U2*-Am30 contain adenosine in 2 adenosines in ‘A<u>A</u>G’ motifs but only adenosine in position 30 within the ‘CA<u>A</u>GUG’ context is *N*^6^-methylated by METTL4 (Figure 2B-2D, S2B). This suggests A in the middle of ‘CA<u>A</u>GUG’ to be METTL4’s target substrate sequence, with a preference for Am over A for adenine-*N*^6^-methylation.

### *In* vivo target sequence preference of METTL4

Since METTL4 catalyzes Am *N*^6^-methylation *in vitro*, we next investigated if METTL4 can also do so *in vivo* by rescuing METTL4 expression in *Mettl4*-KO cells. Similar to all other known m6A and m6Am writers, exogenous METTL4 mainly localizes to the nucleus though in rare cases, exogenous METTL4 was also found in the cytoplasm (Figure 3A, S3A). More importantly, overexpressing METTL4^WT^ rescued the loss of m6Am in *U2 snRNA* of *Mettl4*-KO cells (Figure 3B). On the other hand, overexpressing METTL4^CD^ did not result in the same rescue effect, indicating that METTL4^WT^ is directly catalyzing *N*^6^-methylation of *U2 snRNA* Am *in vivo* (Figure 3B).

**Figure 3.**
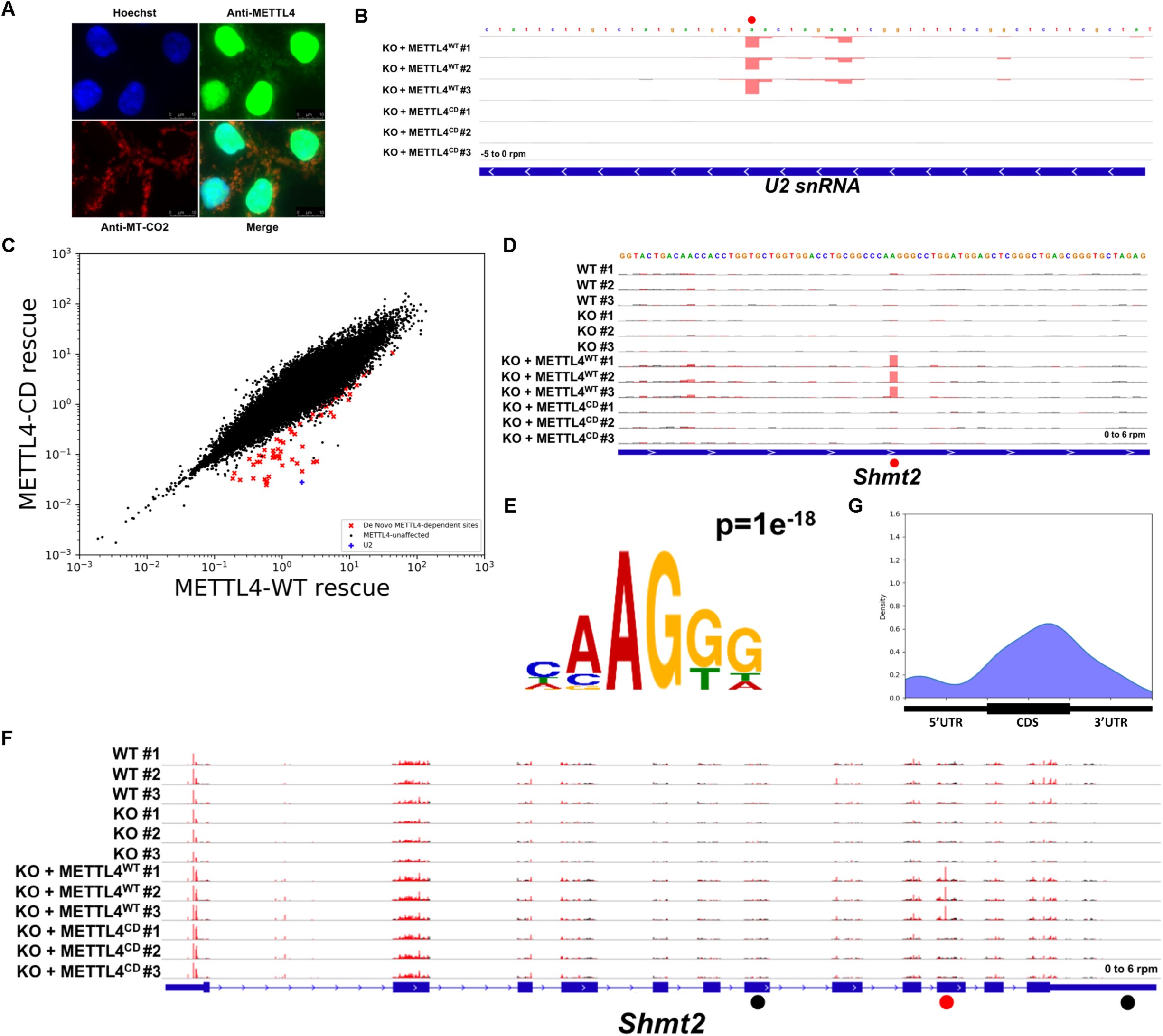
*In* vivo target sequence preference of METTL4. (A) Immunofluorescence of exogenous METTL4 expressed in *Mettl4*-KO cells. Hoechst and anti-MT-CO2 acts as nuclear and mitochondrial markers respectively. (B,D,F) m6ACE (red) and Input (black) read-start counts mapped to various genes. Red dots denote identified *N*^6^-methyladenine positions. Black dots denote ‘HMAGKD’ sites that are not identified as methylated by m6ACE-seq. Sequence corresponds to the same strand as the methylation site. Blue horizontal bar represents transcript. (D) is a magnified version of (F). (C) Scatterplot of average RML of METTL4^WT^-rescue or METTL4^CD^-rescue cells. *De* novo METTL4-dependent are sites with RML reductions of at least log_2_fold of 2 (FDR < 0.1) in METTL4^CD^-rescue compared to METTL4^WT^-rescue cells. (E,G) MEME analysis (E) and metagene distribution profile (G) of METTL4-dependent *de novo* methylation sites.

While we had previously described the preferred RNA target for METTL4 catalysis, we envisioned that treating an extensive variety of RNA sequences with METTL4 will provide a clearer picture of its target sequence preference. Fortunately, the overexpression of METTL4^WT^ in human cells affords such an opportunity: since METTL4^WT^ is capable of methylating *U2 snRNA in vivo*, exposing METTL4^WT^ to the entire human transcriptome allows us to simulate an *in vivo* methylation assay and determine METTL4’s target sequence motif by identifying additional sites of specific RNA methylation. As such, we compared the global methylomes of the METTL4^WT^- and METTL4^CD^-rescue cells to determine if METTL4 also mediates adenosine-*N*^6^-methylation of other RNAs besides *U2 snRNA*. This revealed that overexpressing METTL4^WT^, but not METTL4^CD^, led to the appearance of multiple *N*^6^-methyl-adenine sites in several mRNAs (Figure 3C, Table 2). Since these sites are absent in WT cells, we denote them as *de novo* RNA methylation sites (Figure 3D, S3B). Amongst these *de novo* methylation sites, we identified the dominant motif sequence to be ‘HM<u>A</u>GKD’ (Figure 3E, H=A/C/U, M=A/C, K=G/U, D=A/G/U, <u>A</u> is methylation site). Since the *U2 snRNA* ‘CAm6AmGUG’ motif is a subset of the ‘HM<u>A</u>GKD’, this further validates ‘HM<u>A</u>GKD’ to be METTL4’s target sequence motif.

We next assessed if the presence of a ‘HM<u>A</u>GKD’ sequence is the sole criterion for *de novo* methylation by METTL4^WT^ or if there exists other *cis*-acting elements and/or *trans*-acting factors that guide this process. We first speculated if mRNA expression level is one such factor. However, we found that the same mRNA can possess multiple ‘HMAGKD’ sequences but only have one being *de novo* methylated, thus ruling out mRNA abundance as a key criterion (Figure 3F, S3C). We next considered that METTL4 preferred to methylate Am over A, suggesting that all *de novo* methylated targets initially contained ‘HMAmGKD’. As such, we analyzed Am methylomes mapped using Nm-seq but found no overlap between METTL4^WT^ targeted sites and previously mapped Am sites within the human transcriptome (30). However, we do not completely rule out Am presence as a criterion since Am sites mapped by Nm-seq have yet to be fully validated and there could be multiple false negatives (31). Finally, we assessed if *de novo N*^6^-methylated adenosine exhibited specific positional patterns along the mRNA length. This revealed a slight preference of METTL4^WT^-dependent methylation to be within the CDS (Figure 3G). This suggests that *cis*-acting elements or *trans*-acting factors associated with the CDS help to guide *de novo* methylation by METTL4^WT^.

### METTL4 regulates pre-messenger RNA splicing

While loss of *Mettl4* had no appreciable effect on cell growth rate (Figure S4A), given the role of *U2* in pre-mRNA splicing, we next performed RNA sequencing on WT and *Mettl4*-KO HEK293T cells to search for splicing changes. By applying a stringent criterion to identify the most reliable splicing changes (see Methods), we found a total of 296 splicing events in 165 genes that change upon KO of *Mettl4*, with a majority of cassette exons over differential intron retention, and more alternative 3’ splice sites than 5’ splice sites (Figure 4A). Among the total of 214 affected cassette exons, there are 129 (∼60%) events with enhanced exclusion, and 85 (40%) with increased inclusion in *Mettl4*-KO cells (Figure 4B). A subset of splicing changes map to unannotated splice sites or exons (Figure 4C, S4B). Indeed, we found an excess of unannotated alternative 3’ splice sites over unannotated 5’ splice sites, while most affected cassette exons are annotated.

**Figure 4.**
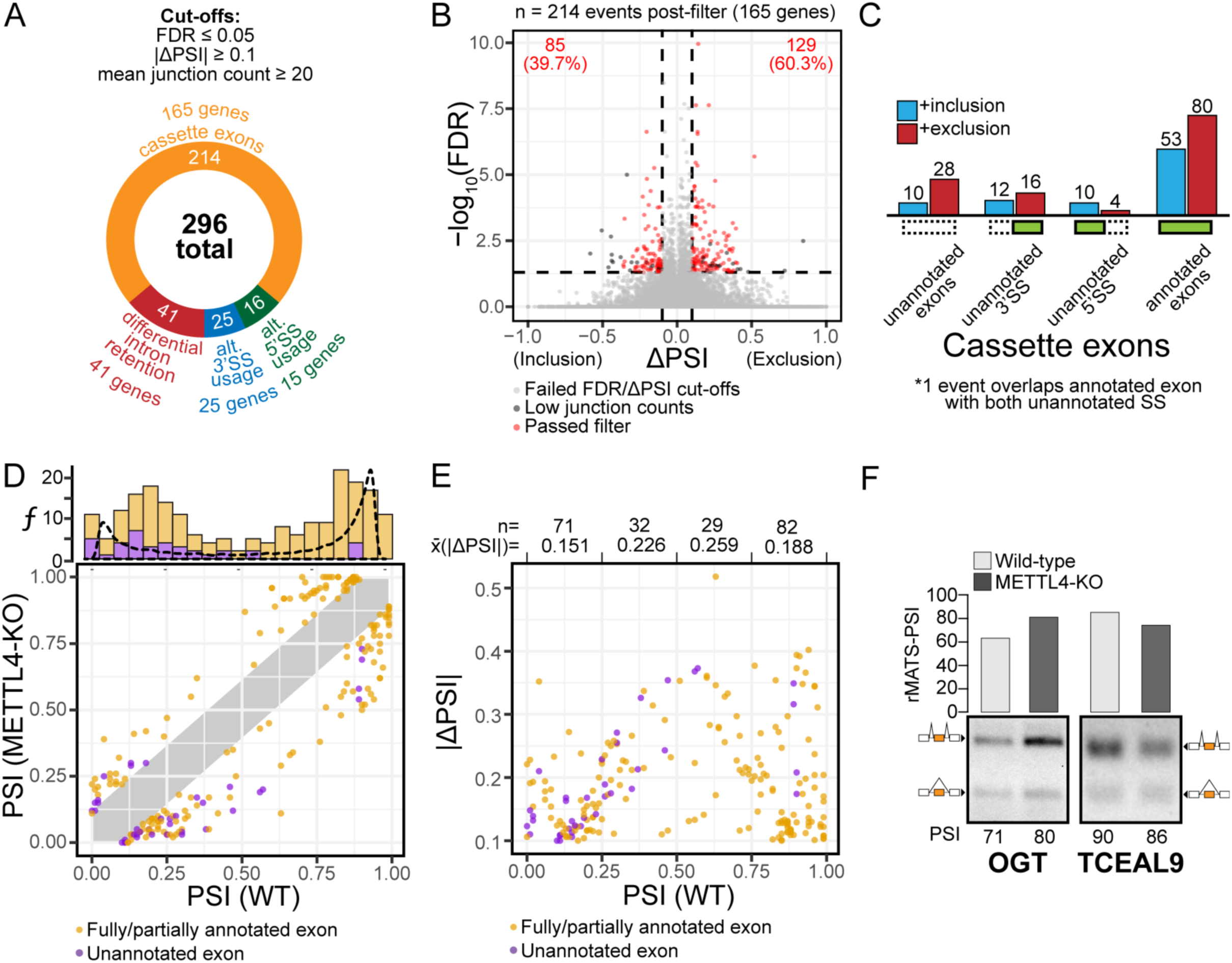
Summary of splicing changes upon Mettl4-KO. (A) Breakdown of the 296 differential alternative splicing events between wild-type and METTL4-KO HEK293T, as reported by rMATS, and with the indicated cutoffs. (B) Volcano plot of differentially spliced cassette exons, with events that passed the filters denoted as red dots. (C) Distribution of the differentially spliced cassette exons based on the directionality of splicing change and annotation status. (D) Plot of initial PSI (WT) against final PSI (*Mettl4*-KO), categorized into annotated exons (orange) and unannotated exons (purple). The histogram represents the relative distribution of initial PSIs with the dotted line representing background initial PSI frequencies. (E) Plot of initial PSI against the absolute changes in PSI values, divided into four equal X-axis intervals and annotated with the mean change in PSI per interval. (F) RT-PCR validations of two differential alternative splicing events for cassette exons in *Ogt* and *Tceal9* genes. The RNA-seq PSI is aligned to the RT-PCR results in agarose gels with its corresponding PSI. The identity of splicing products by exon inclusion or exclusion is schematically shown next to the gels.

Most of the splicing events affected in *Mettl4*-KO cells had either a high or low starting splicing level or PSI (percentage spliced in) while events with intermediate PSI were affected in lower numbers (Figure 4D). As expected, the affected unannotated splicing events tended to have a low starting PSI (Figure 4D, upper chart). As shown before, the magnitude of splicing change (ΔPSI) was the largest for events with intermediate starting PSI (Figure 4E) (32). We show RT-PCR validation of two representative cassette exons for which *Mettl4*-KO increases inclusion such as *Ogt*, or increases exclusion such as *Tceal9* (Figure 4F).

Last, DAVID analysis revealed that the set of genes with differential alternative splicing events upon *Mettl4*-KO is enriched in nuclear processes such as DNA repair and RNA binding or processing (Figure S4C) (33, 34). Hence, the enriched categories might just reflect a preponderance of genes encoding splicing factors or related processes that are themselves regulated by alternative splicing (35).

### Features of splicing events affected by *Mettl4*-KO

Analysis of splice-site strength using the MAXENT algorithm revealed that the annotated cassette exons with increased exclusion in *Mettl4*-KO have weak 3’ splice sites (Figure 5A) (36). The other subcategories in 3’ splice sites and all the 5’ splice-site analyses showed no statistically significant differences. This included the unannotated 3’ splice sites, probably because of the low number of detected splicing events in this group. The exon and intron lengths associated with the splicing events altered in *Mettl4*-KO showed no statistically significant differences either (Figure S4D-S4F).

**Figure 5.**
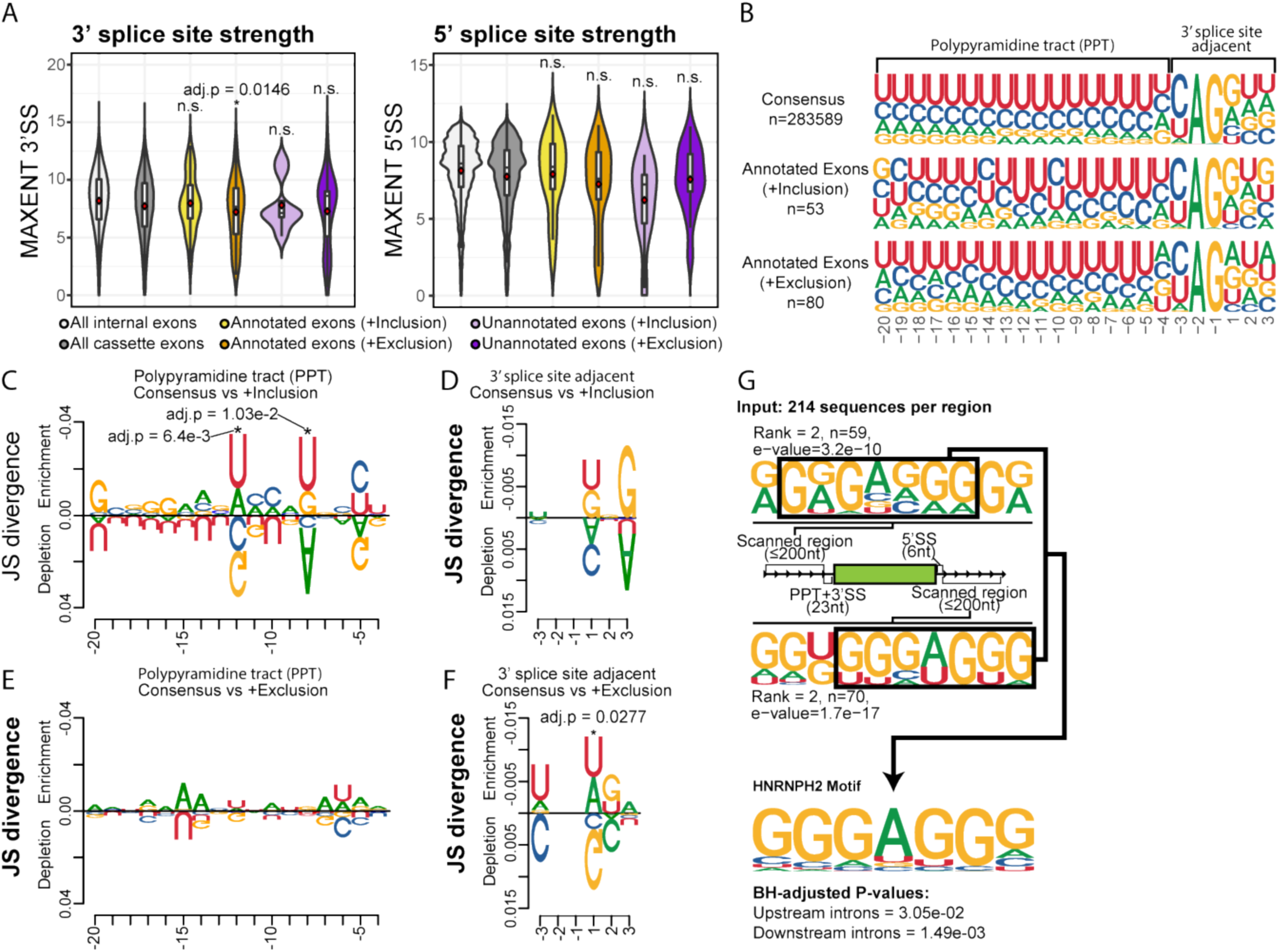
Features of affected METTL4-dependent splicing events. (A) Violin plot of the MAXENT splice-site strength of all human internal exons (n=283589), all human cassette exons (n=63808), annotated exons undergoing increased inclusion (n=53) or exclusion (n=80), and unannotated exons undergoing increased inclusion (n=10) or exclusion (n=28). Mean values are indicated as red diamonds and statistical significance was calculated against all human cassette exons subset. (B) Sequence logos of the consensus 3’ splice-site sequence motif and those of the 133 fully annotated cassette exons that are differentially spliced, categorized by directionality of splicing change. (C-F) Jensen-Shannon Divergence plots comparing the polypyrimidine tract (PPT, C,E) and 3’ splice site adjacent region (D,F) nucleotide compositions of the consensus and experimental 3’ splice site sequences. (G) Sequence logos of enriched sequences on the regions upstream and downstream of differentially spliced cassette exons (n=214) revealed by MEME, that are statistically significant matches for the consensus binding motif of HNRNPH2.

A more detailed look at the affected 3’ splice sites associated with either increased exclusion or inclusion revealed differences in both their polypyrimidine tracts and in the sequence surrounding the AG dinucleotide at the intron-exon junction (Figure 5B). Comparison of the 3’ splice-site motifs by Jensen-Shanon divergence revealed that the polypyrimidine tract motif for increased inclusion events exhibits two positions (−8 and −12) with a significantly higher incidence of uridines than purines, while other upstream positions have a lower frequency of uridines, albeit not reaching significance (Figure 5C). The polypyrimidine tract motif for increased exclusion events did not reveal any position with a significant change (Figure 5D). In turn, the motif around the terminal AG dinucleotide did not show significant differences for increased inclusion events (Figure 5E) yet for increased exclusion, it revealed a significantly reduced frequency of the consensus G at the first exonic position (Figure 5F). Last, motif analyses for 5’ splice sites revealed just a depletion of Cs at position −2 for increased exclusion events (Figure S4G), which might likely be a spurious hit. Overall, these analyses reveal that the 3’ splice sites affected upon *Mettl4*-KO exhibit distinct characteristics potentially related to the absence of m6Am at position 30 of the *U2 snRNA*.

Finally, we searched for enriched motifs in the 200-nucleotide intronic fragments flanking the affected splice sites and exons, and found a very clear GGGAGGG consensus motif on both upstream and downstream introns (Figure 5G). This motif corresponds to the binding site of hnRNP-H2 protein (37). There were a few more enriched motifs including C-rich upstream intronic sequences (Figure S4H). Overall, the METTL4-dependent splicing events exhibit particular 3’ splice-site features possibly linked to *U2 snRNA* modification, yet other mechanisms mediated by RNA Binding Proteins (RBPs) may also apply.

## Discussion

Nucleotide 30 of *U2 snRNA* is an adenosine known to be ribose-2’O-methylated to give Am (28). Recently, the same adenosine was also shown to be *N*^6^-methylated to m6Am but the writer that catalyzes this *N*^6^-methylation was unknown (20). Based on a past phylogenetic prediction of METTL4 as a *N*^6^-methylation writer (29), we knocked out human *Mettl4* and sequenced the resulting loss of RNA methylation, thereby confirming that METTL4 is required for *N*^6^-methylation at adenosine 30 of *U2 snRNA*. We also used nucleoside HPLC-MS/MS to validate the identity of the resultant modification as m6Am. By overexpressing recombinant METTL4, we demonstrated the necessity of METTL4’s catalytic site in its methylation activity and that METTL4 is able to directly methylate *U2 snRNA* both *in vitro* and *in vivo*.

*N*^6^-methylation writers such as METTL3 and PCIF1 have multiple *in vivo* targets and thus by comparing the difference in precise RNA methylomes before and after depleting these writers in cells, we can determine the *in vivo* target RNA preference for each writer (27). METTL4 however, only has 1 clear *in vivo* target, making such an analysis non-trivial. We circumvented this limitation by overexpressing either METTL4^WT^ or METTL4^CD^ in HEK293T cells to simulate an *in vivo* methylation assay. Comparing differences between the RNA methylomes in the 2 overexpressed cells revealed METTL4^WT^-dependent *de novo* methylation sites, allowing us to determine METTL4’s *in vivo* target preference to be ‘HM<u>A</u>GKD’. We note that presence of the ‘HMAGKD’ sequence alone does not guarantee *in vivo de novo* methylation by METTL4. This indicates that other *cis*-acting elements, such as RNA secondary structure or Am presence, or *trans*-acting factors, such as METTL4 co-factors help to guide this process. During the preparation of this manuscript, another study also reported METTL4 as a *U2 snRNA* m6Am writer (38) but due to a lack of m6Am-sequencing, could not conduct the extensive *in vivo* target RNA preference analysis of our study. Therefore, our findings will facilitate further studies on how METTL4 specifically targets *U2 snRNA*.

As opposed to *U2* pseudouridines, which in yeast were shown to affect either *U2* snRNP biogenesis, particle stability, splicing function and even growth phenotypes (39), the role of *U2* m6Am function remains largely uncharacterized. Our transcriptomic analysis revealed a limited yet reliable set of splicing events affected in Mettl4-KO cells, with the typical distribution of subtypes (cassette exons, etc) as compared to most cellular processes or alterations of splicing factors (40). Our meta-analysis of METTL4-dependent differential alternative splicing events strongly suggests that the splicing alterations are governed by 3’-splice site features: (i) A slight overabundance of changes in alternative 3’ splice sites over 5’ splice sites; (ii) an overall weakness of 3’ splice-site strength for cassette exons with increased exclusion in *Mettl4*-KO cells; (iii) specific motif divergences for 3’ splice sites, such as changes in the polypyrimidine tracts for cassette exons with increased inclusion, and decreased consensus nucleotide adjacent to the terminal AG dinucleotide for cassette exons with increased exclusion in KO. These data, together with an overall lack of 5’ splice-site or exon/intron length features, hint that the METTL4-dependent splicing events mainly act via the 3’ splice site, which is directly linked to *U2 snRNA* and likely to its modifications.

In human major spliceosome introns (>99% of all introns), 3’ splice sites are recognized by the pre-formed and stable U2 Auxiliary Factor (U2AF) heterodimer, in which the large U2AF65 subunit binds to the polypyrimidine tract while the small U2AF35 subunit binds a sequence motif around the intron-terminal AG dinucleotide (41-46). Similar to other splicing signals, 3’ splice sites are highly heterogeneous in humans. Hence, the relative contribution of the two subunits to 3’ splice-site recognition is highly variable. This had previously led to the model of AG-dependent versus AG-independent introns, as the latter introns have long polypyrimidine tracts that make U2AF35 binding dispensable for 3’ splice-site recognition (47). U2AF binding to 3’ splice sites together with Splicing Factor 1 (SF1) to the Branch Point Sequence (BPS) then recruits the U2 snRNP in an ATP-dependent manner to displace SF1, such that *U2 snRNA* base pairs to the BPS and bulges out an A to perform the first transesterification step of splicing (48, 49). As the *U2*-30 position is not in the BPS recognition sequence yet 5’-adjacent to it, we hypothesize that the m6Am modification may affect the recruitment of U2 snRNP by U2AF in different manners depending on the pre-mRNA substrate, thereby affecting the overall splicing patterns in a subtle manner. Further studies should aim to identify the mechanistic details of splicing events affected by METTL4 depletion, likely by *in vitro* splicing reactions with reconstituted U2 snRNP particles without and with *U2 snRNA* position 30 m6Am modification, so as to derive the respective *U2* interactomes, degree of *U2* association with pre-mRNAs, and efficiency of spliceosome formation. Furthermore, the enrichment of GGGAGGG motif in both flanking introns of several METLL4-dependent splicing events suggests that some such events might be regulated by the cognate RBP that recognizes this motif, which is hnRNP-H2 (50). This RBP did not undergo any major expression or splicing change at RNA level upon *Mettl4* KO (data not shown), so if this factor accounts for a fraction of splicing events, it is regulated by translation efficiency or post-translational modifications. Further studies should address the connection between METLL4 and hnRNP-H2. Our work here on the discovery of METTL4 as a *U2* m6Am writer provides a framework to explain how METTL4 regulates pre-mRNA splicing, thereby proposing a new pathway for epitranscriptomic modifications to regulate RNA processing.

## Methods

### m6ACE-seq

m6ACE-seq libraries were constructed as previously described (27): Poly(A) RNA was purified using Poly(A)Purist Mag kit (Thermofisher AM1922) according to manufacturer’s instructions, then fragmented to ∼120nt by incubating in RNA fragmentation buffer (Ambion AM8740) for 7.5min at 70°C. Fragmented RNA was treated with 10U T4 PNK (NEB M0201) for 30min at 37°C before adding 1mM ATP and incubating for an additional 30min at 37°C, then purified using Oligo Clean & Concentrator (Zymo D4060). 3’ ligation was performed as described (51, 52), where 200pmol 5’-adenylated,3-dideoxyC DNA adapters (Table 1) were ligated with 400U truncated T4 RNA ligase 2 (NEB M0242) in 1X ATP-free T4 RNA ligase buffer [50mM Tris pH 7.5, 60µg ml^-1^ BSA, 10mM MgCl_2_, 10mM DTT, 12.5% PEG8000] for 2hr at 25°C. Ligated RNA was purified with Ampure XP beads (Beckman Coulter A63881). 200pg of 3’-ligated methylated RNA spike-in (Table 1) was added to 1µg of ligated Poly(A) RNA and the mixture was denatured for 5min at 65°C before incubating for 2min on ice. This denatured RNA mixture was incubated overnight at 4°C with 8µg anti-m6A antibody (Synaptic Systems 202003) in 1X IP buffer [150Mm LiCl, 10mM Tris pH 7.4, 0.1% IGEPAL CA-630 (Sigma I8896)] supplemented with 1U µl^-1^ RNasin Plus (Promega N2611). In parallel, 1.2mg Dynabeads-Protein-A was blocked overnight at 4°C in 1X IP buffer supplemented with 0.5mg ml^-1^ BSA (Sigma A7906). The antibody-RNA mixture was split into 50µl aliquots on ice and crosslinked with 0.15J cm^-2^ 254nm UV radiation six times. The antibody-RNA mixture was recombined and 1% of it was set aside as input-RNA and the remainder (designated as m6ACE-RNA) was mixed with decanted BSA-blocked Dynabeads-Protein-A for 1.5hr at 4°C. Beads bound with crosslinked RNA were then washed with 250µl of the following cold buffers in this order: Wash buffer 1 [1M NaCl, 50mM HEPES-KOH pH 7.4, 1% Triton X-100, 0.1% Sodium Deoxycholate, 2mM EDTA], Wash buffer 2 [0.5M NaCl, 50mM HEPES-KOH pH 7.4, 1% IGEPAL, 0.1% Sodium Deoxycholate, 2mM EDTA], Wash buffer 3 [1% Sodium Deoxycholate, 25mM LiCl, 10mM Tris pH 8, 1% Triton X-100, 2mM EDTA], TE [10mM Tris pH 8, 1mM EDTA] and finally 10mM Tris pH 8. m6ACE RNA was then denatured in 10µl 10mM Tris pH 8 for 5min at 65°C and for 2min on ice. m6ACE RNA was digested with 1U XRN-1 (NEB M0338) in XRN-1 buffer [100mM LiCl, 45mM Tris pH 8, 10mM MgCl_2_, 1mM DTT] and 1U µl^-1^ RNasin Plus shaking at 1krpm for 1hr at 37°C. The m6ACE RNA-bead mixture was then washed with Wash buffer 1, Wash buffer 2, Wash buffer 3, TE and 10mM Tris pH 8. Both input and m6ACE RNAs were eluted in elution buffer [1%SDS, 200mM NaCl, 25mM Tris pH 8, 2mM EDTA, 1mg ml^-1^ Proteinase K (Thermo Scientific EO0491)], shaking at 1krpm for 1.5hr at 50°C. RNAs were ethanol-precipitated and ligated to 5pmol 5’adapters (Table 1) with 10U T4 RNA ligase (Ambion AM2140) supplemented with 12.5% PEG8000 and 2U µl^-1^ RNasin Plus for 16hr at 16°C before being purified with Oligo Clean & Concentrator. 5pmol of reverse transcription primer (Table 1) was annealed (72°C 2min, ice 2min) and reverse transcription was performed with 200U SuperscriptIII (Invitrogen 18080) for 1hr at 50°C, with the reaction stopped by incubating for 15min at 70°C. The cDNA was PCR amplified with Phusion High-fidelity PCR mastermix (NEB M0530) and Truseq PCR primers. Finally, primer-dimer and adapter-dimers were removed with Ampure XP beads before undergoing PE75 sequencing on the Illumina Nextseq platform.

### m6ACE-seq analysis

m6ACE-seq analysis was performed as previously described (27): Fastq sequences were first filtered for a quality score of 20, then trimmed of 5’ and 3’ adapter sequences and poly(A) tails using Cutadapt (53). The 8-mer UMI located at the first 8 nucleotides of read 1 was registered and trimmed. Any complementary UMI sequence in read 2 was also trimmed. Reads were mapped to the methylated spike-in (Table 1) using Bowtie2, or to the hg38 assembly transcriptome (Gencode v28 comprehensive gene annotations) using STAR (54, 55). Aligned pairs that had the same mapping coordinates and UMIs were filtered out as PCR duplicates. Read-start coordinates in hg38-mapped reads that began with an adenosine nucleotide, and had a minimum mean read count of 1 across the triplicate samples were collated. m6A or m6Am sites were identified as read starts that were at least 2-fold enriched in m6ACE libraries than in the corresponding input libraries. This enrichment was calculated using DESeq2 (56) performed on A-only sites across triplicate pairs of m6ACE and corresponding input libraries (FDR<0.1, padj<0.05). Identified sites that were 1-4 nucleotides upstream of another identified significant Rm6AC site or sites found within clustered read-starts were filtered out.

To calculate the RML of each site in each sample: The read-start counts at positions −4 to 0 of each site in the m6ACE library were summed and divided by the read-start counts at positions −51 to 0 of the same site in the input library to give ‘X’. Similarly, the read-start counts at positions −4 to 0 of the spike-in m6A site in the m6ACE library were summed and divided by the read-start counts at positions −21 to 0 of the same spike-in m6A site in the input library to give ‘Y’. X was normalized to Y to give RML. RML values of each site was averaged across triplicates for each sample condition. A site was denoted as differentially methylated between METTL4^WT^-rescue and METTL4^CD^-rescue if the average RML differs between sample conditions with a log_2_fold-change cutoff of 2.0, as well as a one-tailed Student’s T-test p-value cutoff of <0.041 (false discovery rate, FDR < 0.1). Consensus motif analysis was performed using Meme-chip (37). Metagene analysis was performed using MetaPlotR (57).

### RNAseq

10µg of RNA was treated with RQ1 RNase-Free DNase (Promega M6101) as per manufacturer’s protocol. 5µg of samples were then depleted of rRNA using the ribozero kit (Illumina MRZH11124). 50ng of each rRNA-depleted samples were then used for construct RNA-seq libraries using the ScriptSeq v2 RNASeq Library Preparation Kit (Illumina SSV21124). Libraries were sequenced on an Illumina PE151 platform.

### RNAseq Splicing Analysis

WT and *Mettl4*-KO HEK293T RNAseq libraries were first processed using CutAdapt to remove adaptors and to trim all reads to a uniform length of 101bp for compatibility with downstream analysis steps. Read alignment to the genome was then performed using STAR (2.7.0a) in two-pass mode using the hg38 genome assembly and GENCODE v28 gene annotations (55).

Alternative splicing differences were analysed by an in-house pipeline using replicate MATS (rMATS 3.2.5) and filtered based on cut-offs of FDR ≤ 0.05, an absolute change in percent spliced in (|ΔPSI|) ≥ 0.1 (58, 59). In addition, events with mean sample junction counts (IC_Sample + SC_Sample) < 20 were filtered out to reduce false positives arising from transcript dropout in low abundance isoforms. The validation of several differential alternative splicing events were performed using an RT-PCR assay (36, 59).

Additional exon-level features were added to the data using databases constructed from the UCSC Table Browser, as well as in-house R scripts. Splice-site strengths were calculated using MaxEntScan (36). Comparisons were made between the exon features of the differentially spliced cassette exons and a transcriptome-wide background based on all known human internal exons. Cut-offs of 500nt and 30,000nt were applied on the exon and intron length datasets respectively to reduce data skewing caused by extreme outliers. Splice site strength values below 0 were also discarded from the analysis for similar reasons. The statistical significance of feature differences were calculated using Welch’s t-test and p-values were adjusted using Bonferroni correction when necessary.

The Jensen-Shannon Divergences of splice site sequence subsets and associated tests for statistical significance were calculated using DiffLogo (60). All other sequence logos were generated using ggseqlogo (61). The cassette exon sequences, as well as up to 200nt stretches up- and downstream of the exon (excluding the splice sites), were scanned for strand-specific 6-10nt motifs using MEME (37). The sequences generated by MEME were then fed into TOMTOM to search for RNA binding proteins with similar binding motifs. Functional annotation of the genes containing differentially spliced cassette exons was performed using DAVID (33, 34).

### RT-PCR assay

1µg of total RNA was primed with random hexamers and reverse transcribed with SuperscriptIII (ThermoFisher Scientific 18080044) as per manufacturer’s protocol. 1µl of 5U/ul RNAse H (ThermoFisher Scientific EN0201) was then added to each reaction and incubated at 37°C for 20min. DNA was used for PCR amplified for 35 cycles using Taq polymerase (NEB M0273) and respective primers (Table 1) before being assessed via agarose gel electrophoresis.

## Acknowledgements

We thank the Lezhava laboratory for assistance with sequencing. This work was supported by the Biomedical Research Council (A*STAR) Young Investigator Grant 1610151037 (W.S.S.G.), the Genome Institute of Singapore Independent Fellowship (W.S.S.G.), as well as Singapore’s Ministry of Education Academic Research Fund Tier 2 grant MOE2016-T2-2-104(S) (X.R.).

## Author Contributions

W.S.S.G. conceived the study. Y.T.G. and C.W.Q.K. performed the experiments. D.Y.S. and W.S.S.G. performed bioinformatics analysis. X.R. and W.S.S.G. supervised the experiments and wrote the manuscript with input from all authors.

