## Supplementary figures and images for "METTL4 catalyzes m6Am methylation in *U2 snRNA* to regulate pre-mRNA splicing"

### Supplemental Data 1

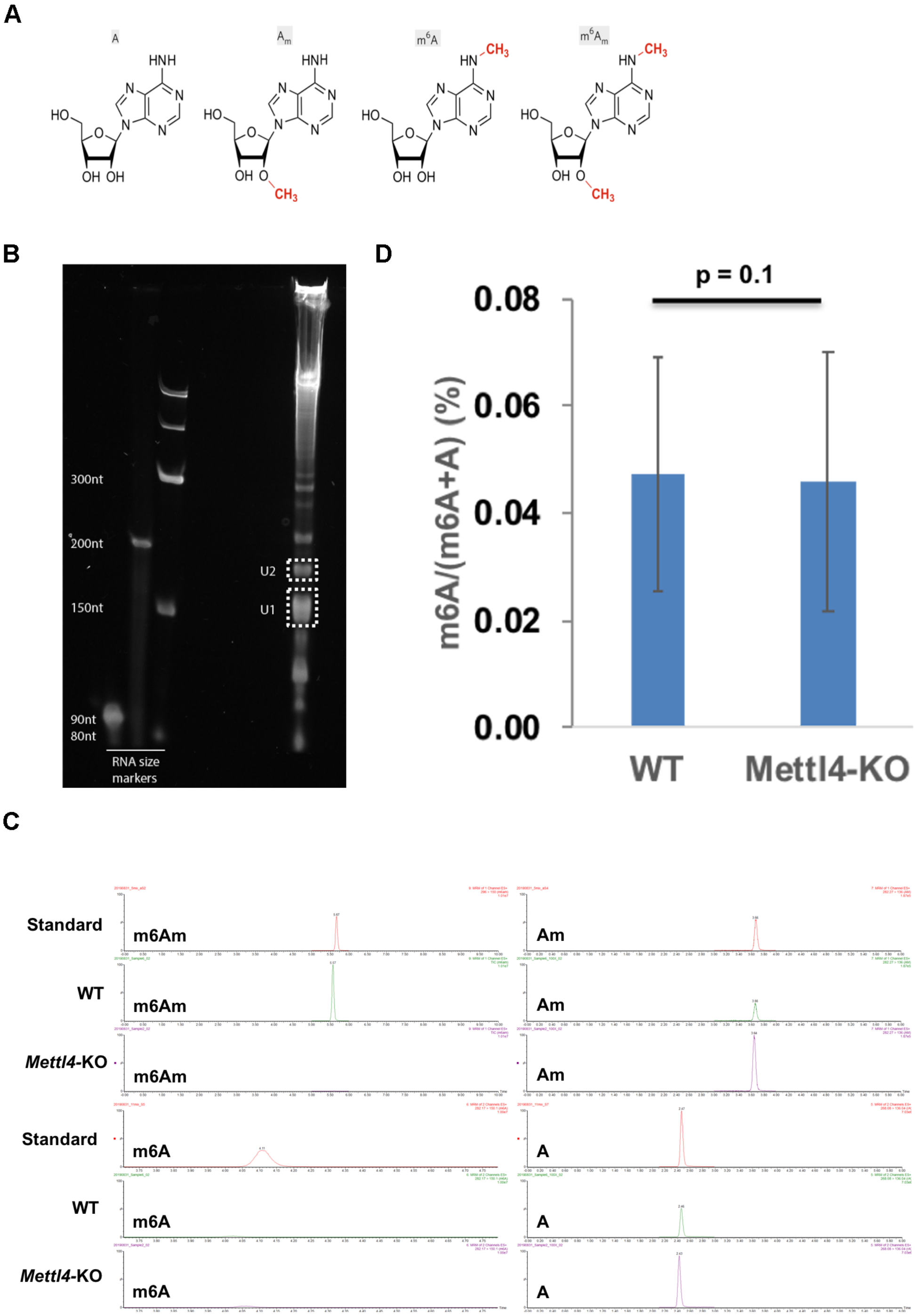

### Supplemental Data 2

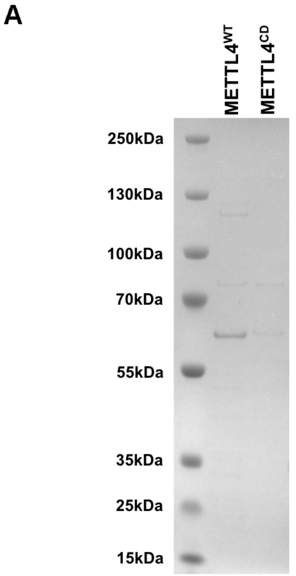

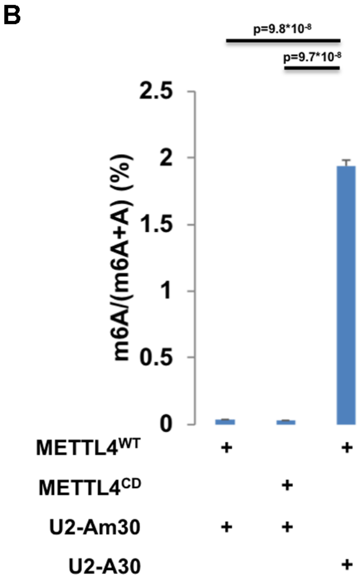

### Supplemental Data 3

**FIGURE S3**

**A**

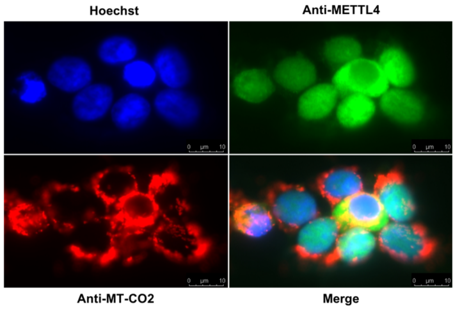


**B**

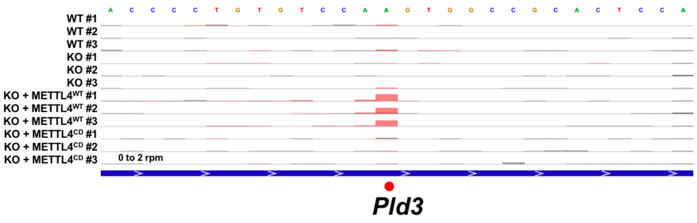


**C**

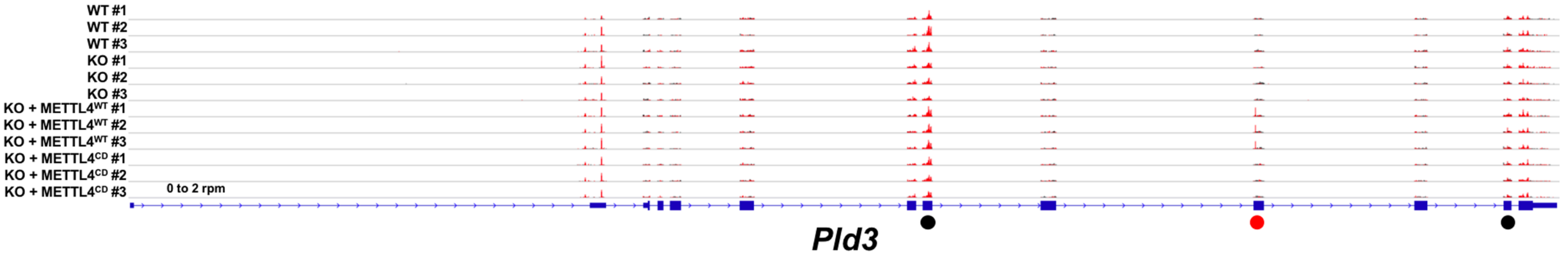

### Supplemental Data 4

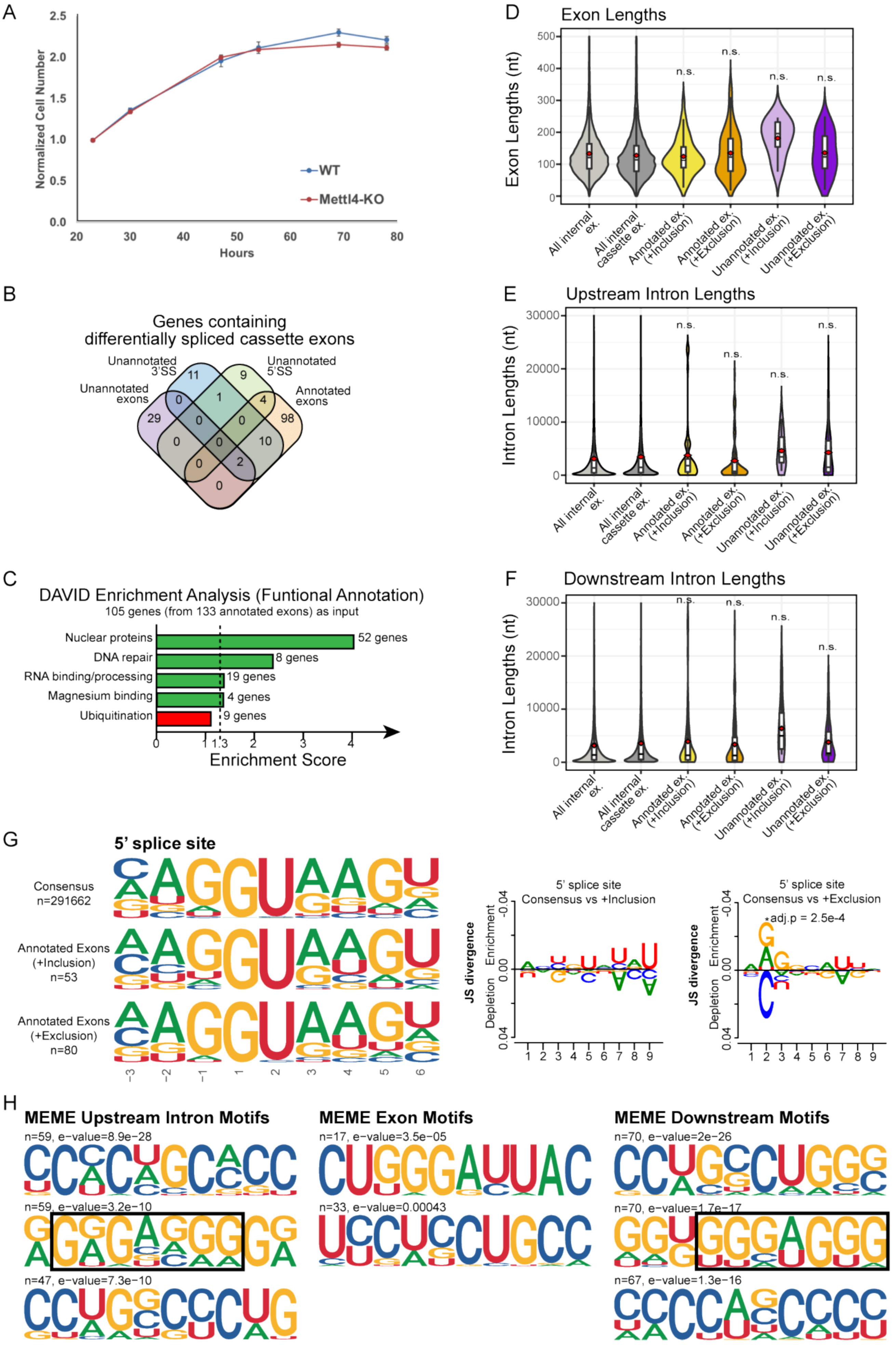
