## Supplemental Data 5 for "METTL4 catalyzes m6Am methylation in *U2 snRNA* to regulate pre-mRNA splicing"

**Goh *et al.* Supplementary Information**

**Supplementary Figure Legends**

**Figure S1. METTL4 mediates m6Am modification in U2 snRNA.**

(A) Chemical structures of A, Am, m6A and m6Am.

(B) Nuclear-enriched RNA separated on a TBE-urea gel. Dotted rectangles denote cut sites for isolation of various snRNAs based on their sizes.

(C) HPLC retention time chromatograms for various nucleosides standards and of WT and *Mettl4*-KO U2 snRNA samples.

(D) Nucleoside HPLC-MS/MS of m6A as a percentage of total m6A and A in U2 snRNA purified from WT versus *Mettl4*-KO RNA. 1-tailed Student's T-test p-value for triplicates is shown.

**Figure S2. METTL4 directly catalyzes m6Am formation *in vitro***

(A) SDS-PAGE gel image of purified C-terminal 3X-FLAG-tagged METTL4$^{WT}$ and METTL4$^{CD}$. Expected size of purified protein is 57kDa.

(B) Nucleoside HPLC-MS/MS of m6A as a percentage of total m6A and A in RNA substrates *in vitro* methylated by METTL4$^{WT}$ or METTL4$^{CD}$. Displayed are average and standard deviation error of triplicates. 1-tailed Student's T-test p-value is shown.

**Figure S3. *In* vivo target sequence preference of METTL4**

(A) Immunofluorescence of exogenous METTL4 expressed in *Mettl4*-KO cells. Hoechst and anti-MT-CO2 acts as nuclear and mitochondrial markers.

(B,C) m6ACE (red) and Input (black) read-start counts mapped to various genes. Red dots denote identified $N^6$-methyladenine positions. Black dots denote 'HMAGKD' sites that are note identified as methylated by m6ACE-seq. Sequence corresponds to the same strand as the methylation site. Blue horizontal bar represents transcript. (B) is a magnified version of (C).

**Figure S4. Features of affected METTL4-dependent splicing events**

(A) Cell numbers of WT versus *Mettl4*-KO cells grown over time and measured using Alamar blue assay. Cell numbers are normalized to that of the first time point.

(B) Distribution of genes containing differentially spliced cassette exon events, as subsets of those with unannotated exons (n = 31), exons with unannotated 3' splice site (n = 24), exons with unannotated 5' splice site (n = 14) and fully annotated exons (n = 114).

(C) Functional enrichment of the 105 genes from the 'fully annotated exon' subset using DAVID. 9 non-coding genes were excluded from the analysis.

(D-F) Distribution of exon lengths, upstream intron lengths and downstream intron lengths of all human internal exons (n=283589), all human cassette exons (n=63808), annotated exons undergoing increased inclusion (n=53) or exclusion (n=80), and unannotated exons undergoing increased inclusion (n=10) or exclusion (n=28). Mean values are indicated as red diamonds and statistical significance was calculated against the all human cassette exons subset.

(G) Sequence logos of the consensus 5' splice-site sequence motif and those of the 133 fully annotated cassette exons that are differentially spliced, categorized by directionality of splicing change.

(H) Sequence logos of the top 3 enriched sequences on the regions upstream and downstream of differentially spliced cassette exons (n=214) revealed by MEME, as well as the two significantly enriched sequences within the exon body itself.

### Methods

#### Tissue Culture

ATCC HEK293T CRL-3216 cells were cultivated in a sterile 5% $CO_2$ incubator at 37$^{o}$C, and in DMEM supplemented with 10% FBS and 1% penicillin/streptomycin. Cells within passage 3-20 were used for experiments. HEK293T cells were regularly subjected to MycoAlert Plus Mycoplasma kit (Lonza LT07) to verify that they were mycoplasma-free.

##### Generation of Knockout Cell Lines using CRISPR-cas9

HEK293T gene deletions were performed as described. Briefly, guide RNA sequences corresponding to a region around the start codon of the gene of interest were designed using CRISPOR, then cloned into pSpCas9 BB-2A-puro (Addgene 62988) plasmids. Pairs of guide RNAs (Table 1) were designed to induce either frameshift mutation close to the 5'end of the gene or to delete the start codon. HEK293T cells were plated in 12-well plates at $2*10^5$ cells per well in regular growth media but without antibiotics. 16-24hr later, cells were transfected with 500ng of each of the pair of guide RNA-expressing plasmid via Lipofectamine 2000 (Thermofisher 11668). 24hr post-transfection, successfully transfected cells were selected via survival under a 72hr treatment with 2µg $ml^{-1}$ puromycin. Puromycin-resistant cells were expanded for monoclonal dilution to select for monoclones that present desired gene deletions. The knockout mutations in these monoclones were further verified by loss of protein of interest via Western blotting.

#### RNA isolation

Total RNA was isolated from adherent HEK293T using Trizol-LS (Ambion 10296) according to manufacturer's instructions and quantified using the Qubit RNA HS assay (ThermoFisher Q32855).

**Total protein isolation**

Trypsinized HEK293T cells were washed twice with ice-cold PBS. Washed cells or intact nuclei were lysed in RIPA buffer [150mM NaCl, 1% NP-40, 0.5% sodium deoxycholate, 0.1% SDS, 50mM Tris pH 8, 1X Complete Mini EDTA-free protease inhibitor] by tumbling for 30mins at 4$^oC$. Lysate was clarified by centrifuging at 16,000g for 30mins at 4$^oC$ and protein concentration was quantified Pierce BCA protein assay kit (Thermo Scientific 23225).

**Western Blotting**

Whole cells or cell fractionations were lysed in RIPA buffer [150mM NaCl, 1% NP-40, 0.5% sodium deoxycholate, 0.1% SDS, 50mM Tris pH 8, 1X Complete EDTA-free protease inhibitor cocktail] and protein concentration was quantified via BCA assay. Whole cell or mitochondrial lysates were then diluted in 1x Laemmli buffer (Biorad 1610747) supplemented with 2-mercaptoethanol, before being denatured for 10min at 95$^oC$. ~30µg lysate was separated on a 10% SDS-PAGE gel in separation buffer [25mM Tris pH 8.3, 192mM glycine, 0.1% SDS] and transferred onto a nitrocellulose membrane via wet transfer in cold transfer buffer [25mM Tris pH 8.3, 192mM glycine, 20% methanol]. The membrane was rinsed with water then blocked with Odyssey blocking buffer (Licor 927) for 1hr at room temperature. Membrane was stained for 16hr at 4$^oC$ overnight with antibody dilution solution [0.1% Tween-20 in Odyssey blocking buffer] containing primary antibodies. Membrane was rinsed then washed thrice with PBS-T [1X PBS, 0.1% Tween20], before being stained with secondary antibodies. Membrane was rinsed then washed thrice with PBS-T and once with PBS before being imaged on a Licor Odyssey CLx imaging system. The following antibodies and dilution scales were used: 1µg $ml^{-1}$ Mouse anti-actin (Santa Cruz sc-8432); 250x diluted rabbit anti-METTL4 (Sigma HPA040061); 200x diluted mouse anti-HSP60 (Abcam ab110312); 10,000x diluted IRDye 680RD goat anti-mouse IgG H+L (Licor 68070) and 10,000x diluted IRDye 800CW goat anti-rabbit IgG H+L (Licor 32211).

**Immunofluorescence**

Cells were seeded on poly-D-lysine coated coverslips (Neuvitro GG-14-PDL) in 24-well plates 24-48hr before fixing. At ~70% confluency, cells were washed in PBS once and fixed in 4% formaldehyde (diluted in PBS, Thermo 28906) for 10mins. The cells were rinsed and washed thrice using cold PBS. All washes steps were done in 0.5mL volume with shaking at 50rpm for 5mins at room temperature unless otherwise stated. Permeabilization of cell membranes was done using 1% Triton X-100 (diluted in PBS) for 10mins. The cells were rinsed and washed thrice using PBS-T [PBS with 0.1% Tween 20]. Blocking was then done using PBS-T with 10% goat serum (Sigma G9023) for 1hr at RTP. Primary antibodies used are rabbit anti-METTL4 (HPA040061, 1:150), mouse anti-FLAG (Sigma F1804, 1:1,000), and were diluted in antibody dilution buffer [PBS-T with 1% goat serum]. The blocking solution was aspirated and the diluted primary antibody added directly to coverslip before incubating overnight in a 4$^{o}$C humid chamber. After which the cells were rinsed and washed thrice using PBS-T. Fluorescent secondary antibodies (Invitrogen A11019 & A11070) were also diluted in antibody dilution buffer and added directly to the coverslip before incubation in the dark for 1hr at room temperature. The cells were rinsed and washed thrice using DPBS-T in the dark. Hoechst solution was prepared by diluting Hoechst 33342 (Invitrogen H3570) in DPBS. 2µg $ml^{-1}$ of Hoechst solution was used for nuclear staining in the dark for 5mins. The cells were rinsed and washed thrice in DPBS in the dark for 10mins each. The coverslips were then placed onto a drop of Prolong Diamond Antifade Mountant (Thermo P36970) on a glass slide, cured overnight and sealed with nail polish. Images were taken with a Leica DMi8 microscope.

**Generation of overexpression cell lines**

Full-length WT *Mettl4* cDNA or the catalytically-dead form with D287A/W290A mutations were cloned upstream of a 3x-FLAG tag in the p3xFLAG-CMV14 vector. Overexpression cell lines were generated by transfecting respective cloned plasmids into *Mettl4*-KO HEK293T cells with Lipofectamine 2000 reagent according to manufacturer's instructions. Cells were passaged after 24hrs and expanded for an additional 48hrs for RNA extraction, protein extraction or immunofluorescence.

**Recombinant protein purification**

Overexpression cell lines within the *Mettl4*-KO background were trypsinized and $2*10^{7}$ cells were used per FLAG-tag purification. Cells were washed with cold PBS twice and resuspended in 1ml of cold lysis buffer [150mM NaCl, 50mM Tris pH7.4, 1mM EDTA, 0.1% IGEPAL, 1x protease inhibitor, 1x phosphatase inhibitor]. Cells were allowed to lyse on ice for 30mins, with 10min-interval pipette mixing. SigmaPrep spin columns (Sigma MC1000) were equilibrated with 0.5ml of equilibration butter [150mM NaCl, 50mM Tris pH.4, 1mM EDTA, 0.1% IGEPAL], followed by addition of 40ul anti-FLAG M2 agarose resin. The resin was washed a total of 3 times in 0.5ml of equilibration buffer by centrifugation at 1,000g for 5mins at 4$^{o}$C. Cell lysates were clarified at 13000rpm for 10mins at 4$^{o}$C and the supernatants was collected as the protein lysate. Protein lysate was added to the prepared resin and mixed for 2hr at 4$^{o}$C. Bound proteins were washed thrice with 0.5ml of wash buffer [150mM NaCl, 50mM Tris pH7.4, 1mM EDTA, 0.1% IGEPAL, 1x protease inhibitor] by mixing for 10min at 4$^{o}$C and centrifuging at 1000g for 2mins at 4$^{o}$C.

Bound proteins were eluted with 100μl of elution buffer [150mM NaCl, 50mM Tris pH7.4, 1mM EDTA, 1x protease inhibitor, 0.15μg/μl 3xFLAG peptide (Sigma F4799)] by mixing for 30mins at 4°C, followed by centrifuging at 1000g for 2mins at 4°C to collect the eluate. A total of 3 elutions was collected. Glycerol was added to individual elutions to obtain 20% final glycerol concentration.

**In vitro methylation assay (IVM)**

U2-A30 and U2-Am30 RNA oligonucleotides (Table 1) were diluted to 50μM in water and linearized by incubating at 65°C for 5mins, before immediately placing on ice. IVM reaction was performed in a 25μl volume buffer with a final concentration of 50mM HEPES-KOH pH7.4, 50mM NaCl, 0.1mM $MgCl_2$, 25μM SAM, 1mM DTT, 1U/μl RNasin, 2μM linearized RNA and the respective recombinant protein. The IVM reaction was incubated for 2hrs at 37°C and stopped by adding Trizol-LS. RNA was purified according to manufacturer's protocol, then subjected to dot blotting or nucleoside HPLC-MS/MS.

**Dot blotting**

Dot blotting was performed using the BioRad dot blot apparatus. Hybond N+ membrane (GE Healthcare) was washed with water before addition of 100ng of RNA samples per well. RNA samples were kept on ice and denatured with a final concentration of 1x ice-cold denaturing buffer [10mM NaOH, 1mM EDTA] just before adding to the wells. After the addition of RNA samples, the membrane was washed with 1 round of ice cold 1x denaturing buffer through the wells. Membrane was collected and air-dried for 5 minutes. RNA was then crosslinked to membrane with 0.12J 254nm UV radiation twice with a 30seconds rest in between. The membrane was blocked with Odyssey blocking buffer (Licor 927) for 1hr at room temperature. Membrane was stained for 16hr at 4°C overnight with antibody dilution solution [0.1% Tween-20 in Odyssey blocking buffer] containing 1ug/ml of rabbit anti m6A polyclonal antibody (Synaptic Systems 202003). Membrane was rinsed then washed thrice with PBS-T [1X PBS, 0.1% Tween20], before being stained with secondary antibodies. Membrane was rinsed then washed thrice with PBS-T and once with PBS before being imaged on a Licor Odyssey CLx imaging system.

**snRNA Isolation**

Nuclear-enriched RNA was resolved on a 6% TBE-urea gel then stained with SYBR-gold (Invitrogen S11494). U1/5.8s-rRNA and U2 snRNAs were purified by cutting out the 164nt and 191nt bands respectively, using the low-range ssRNA ladder as a size marker (NEB N0364). snRNAs were gel eluted twice in elution buffer [0.4M NaCl, 10mM Tris pH 7.5, 1mM EDTA pH 8] at 16°C overnight with shaking at 2,000rpm. RNA was then precipitated with equal volume isopropanol and 20μg glycoblue

(Thermofisher AM9515) and washed with 70% ethanol before the pellet was dissolved in water. snRNAs were used for either dot blotting or nucleoside HPLC-MS/MS.

**Nucleoside HPLC-MS/MS**

100ng RNA was subjected to Nuclease P1 treatment with 0.2mM ZnCl2, 20mM NH4OAc, pH5.3 and 0.033U/µl Nuclease P1 (Sigma N8630) for 2 hours at 42$^{o}$C. Samples were then incubated with a final concentration of 100mM NH4HCO3, 0.025mU/ul phosphodiesterase (Sigma P3243-1VL) and 0.025U/µl E.coli alkaline phosphatase (Sigma P7640) for another 2 hours at 37$^{o}$C then heat inactivated for 5min at 65$^{o}$C. RNA was then stored at -20$^{o}$C until nucleoside HPLC-MS/MS.

Nucleoside UHPLC-MS/MS was performed at the Singapore Phenome Centre as described: for reverse phase liquid chromatograph, a HSS T3 (1.8µm; 2.1x100mm) column was used with the following parameters. Mobile phase A: water + 0.1% formic acid; mobile phase B: acetonitrile + 0.1% formic acid; flow rate: 0.3ml min$^{-1}$; column temperature: 40$^{o}$C; sample temperature: 4$^{o}$C; injection volume: 5µl; sample loop: 5µl. Elution gradient condition was set as 0min 2%B, 0.5min 2%B, 6min 8%B, 6.5min 8%B, 6.6min 2%B, 8min 2%B. Am and m6Am eluted with the retention times of 3.64min and 5.64min respectively. Tandem mass spectrometry was performed using a Xevo TQ-S, Waters machine with the following parameters. Ion mode: ESI positive; Acquisition model: MRM; capillary voltage: 3.2kV; desolvation temperature: 400$^{o}$C; Con gas flow: 150L h$^{-1}$; desolvation gas flow: 800L h$^{-1}$; source temperature: 150$^{o}$C; collision energy: 16eV.

A, m6A, Am and m6Am were detected by monitoring mass transitions of m z$^{-1}$ = 268.08 → 136.04, m z$^{-1}$ = 282.17 → 150.10, m z$^{-1}$ = 282.27 → 136.0 and m z$^{-1}$ = 296.0 → 150.0 respectively. All nucleosides were quantified based on a linear calibration curve generated using A (Sigma A9251) m6A (Berry & Associates PR3732), Am (Berry & Associates PR3734) or m6Am (Carbosynth NM157470) nucleoside standards. All measurements were performed in technical triplicates.

**Cell growth Assay**

Cells were seeded in 100µl media per well (96-well format) in triplicates at 2500 cells per well. After 24hrs, 10ul of the 10% Alamar blue was added to all wells except for wells used for measuring background intensities, where 10ul of media was added to those wells instead. Promega GloMax Discover plate reader was used to measure the absorbance. The absorbance readings of the wells were measured at 560nM and 600nM wavelength and normalized cell numbers were calculated according to manufacturer’s instructions.
